# Different control strategies drive interlimb differences in performance and adaptation during reaching movements in novel dynamics

**DOI:** 10.1101/2022.11.11.516159

**Authors:** David Córdova Bulens, Tyler Cluff, Laurent Blondeau, Robert T. Moore, Philippe Lefèvre, Frédéric Crevecoeur

## Abstract

Humans exhibit lateralization such that most individuals typically show a preference for using one arm over the other for a range of movement tasks. The computational aspects of movement control leading to these differences in skill are not yet understood. It has been hypothesized that the dominant and non-dominant arms differ in terms of the use of predictive or impedance control mechanisms. However, previous studies present confounding factors that prevented clear conclusions: either the performances were compared across two different groups, or in a design in which asymmetrical transfer between limbs could take place. To address these concerns, we studied a reach adaptation task during which healthy volunteers performed movements with their right and left arms in random order. We performed two experiments. Experiment 1 (18 participants) focused on adaptation to the presence of a perturbing force field and Experiment 2 (12 participants) focused on rapid adaptations in feedback responses. The randomization of the left and right arm led to simultaneous adaptation, allowing us to study lateralization in single individuals with symmetrical and minimal transfer between limbs. This design revealed that participants were able to adapt control of both arms, and that adaptation was greater in the dominant arm than in the non-dominant. We also observed that the non-dominant arm showed a different control strategy compatible with robust control when adapting to the force field perturbation. EMG data showed that these differences in control were not caused by differences in co-contraction across the arms. Thus, instead of assuming differences in predictive or reactive control schemes, our data show that in the context of optimal control, both arms can adapt, and that the non-dominant arm uses a more robust, model-free strategy likely to compensate for less accurate internal representations of movement dynamics.

**Significance statement:** We studied a reach adaptation task during which volunteers performed the task with their right and left arm randomly. The randomization of the arms allowed us to study lateralization in single individuals with symmetrical and minimal transfer between limbs. We observed a slightly greater adaptation of the dominant arm in the force applied to counter the perturbation. Moreover, the non-dominant arm showed a more robust control strategy when adapting to the force field perturbation, which enabled similar deviations despite faster movements. These interlimb differences were not caused by differences in co-contraction across the two arms. Our results suggest that both arms can adapt to the presence of a force field but the non-dominant arm uses a more robust, model-free strategy.

## Introduction

Handedness is a prominent feature of human motor behaviour. Indeed, most humans have a natural tendency for using one arm, i.e. the dominant arm, when performing a wide range of movement tasks. Studies have revealed an advantage of the dominant arm in the control of limb dynamics (Bagesteiro and Sainburg, 2002; Sainburg and Kalakanis, 2000) and an advantage of the nondominant arm during tasks that required load compensation (Bagesteiro and Sainburg, 2003). From these results, Sainburg (2002) suggested a specialized role for each arm, such that differences in performance across arms arise from differences in the use of predictive and impedance control strategies. More precisely, this hypothesis suggests that both arms are controlled using a mix of predictive and impedance control mechanisms, with asymmetries in performance arising because the dominant arm relies more heavily on predictive control while the non-dominant arm relies more heavily on impedance control (Yadav and Sainburg, 2014b).

Although attractive, this hypothesis implies that participants can modulate the mechanical impedance of their limb to modify their behaviour, presumably through the combined activation of agonist-antagonist pairs of muscles acting on each joint (E Burdet et al., 2001; Hogan, 1985). However, the intrinsic impedance of muscles is quite low at spontaneous levels of activation, and the presence of co-contraction mostly impacts the gain of the stretch reflex (Crevecoeur and Scott, 2014a; Pruszynski et al., 2009). Furthermore, what is often referred to as limb stiffness includes the contribution of reflexes and early voluntary responses (up to ~300ms, (Burdet et al., 2000)), which are known to largely depend on neural feedback processes (Scott, 2016). Thus, it is unclear if the modulation of limb impedance is sufficient to explain differences across dominant and non-dominant arms in reaching adaptation, and the possible involvement of different feedback control strategies has not been investigated.

Regarding the role of feedback, recent studies reported very small changes in background EMG activity across limbs in perturbation tasks (Maurus et al., 2021; Walker and Perreault, 2015). Moreover, it has been suggested that humans tend to use a more robust control strategy that aims to minimize the impact of a ‘worst-case’ perturbation on the system (Basar and Bernhard, 1995), and relies on an increase in the control gains when facing unpredictable force fields during reaching (Crevecoeur et al., 2019). In the context of reaching movements, this increase in the control gains leads to a greater movement speed and reduced deviation from a straight line when disturbed by mechanical perturbations. Altogether these observations warrant a careful re-examination of the neurophysiological basis of inter-limb differences in the control of reaching movements.

However, recent results have shown that the two arms can develop feedforward adaptation equally well (Reuter et al., 2016; Stockinger et al., 2015), with no significant differences observed in the improvement of kinematic errors between the two arms. Here, we investigated the possibility that, while both arms can develop similar levels of feedforward adaptation, there is a difference in the underlying control strategies being used to perform reaching movements with each arm. To test this, we measured participants’ performance as they learned to move a robotic handle in a force field (Experiment 1) following a standard adaptation paradigm. We also measured participants’ adaptation of feedback responses to force fields applied randomly (Experiment 2). Importantly, in both cases, trials in baseline or force field environments were performed with the left or right arms in random order to minimize the potential transfer of adaptation between limbs that could impact differences in behaviour. We observed that performance improved in both arms with comparable rates. Both arms showed a transient increase in reaching movement speed across trials followed by a decrease in speed that remained above baseline trials, which suggests they relied on a more robust control strategy to counter the presence of the force field. The data indicated that, during the late adaptation phase, the dominant arm of participants showed a slightly better adaptation to the perturbation while the non-dominant arm maintained a more robust strategy across trials. The results are interpreted as the expression of differences in the quality of internal models across the dominant and non-dominant arms.

## Materials and methods

### Participants

Twelve healthy participants (age: 22±1yrs, mean± standard deviation across participants), self-identified as right-handed, participated in Experiment 1. Eighteen participants (age: 25±3yrs), self-identified as right-handed, participated in Experiment 2. All participants provided written informed consent before participating in this study. The volunteers had no known neurological disorders and were naïve to the purpose of the experiment. The experimental procedures were approved by the local ethics committee at Université catholique de Louvain.

### Behavioral task

Both tasks shared the same experimental procedure and only differed in the frequency and orientation of the mechanical perturbations applied during movements. Participants held the handles of two robotic arms (Kinarm, Kingston, ON, Canada), one in each hand. Each handle was equipped with a force sensor (Mini-40 F/T sensors, ATI Industrial Automation, NC, USA). Two-dimensional position, velocity, and force at the handle of each robotic arm were sampled at 1kHz. Direct vision of the limbs and robotic handles was blocked throughout the experiments, but two hand-aligned cursors were always visible. Participants were instructed to position their left and right hands in one of two home visual targets (a filled circle with a 0.6cm radius) representing the starting position of reaching movements. These home targets were located on the right or left side of the workspace equally distant from the midline, such that the position of the home targets was naturally associated with the right or left arm (Figure 1A). Participants first waited in the home target for a random period of 2 to 4s. An empty goal target was presented in front of the corresponding home target (and arm), thereby cueing the arm that was to perform the upcoming trial. After the random wait time, the goal target was filled in instantaneously giving them the go signal. Participants were asked to reach the target between 600 and 800ms after the go signal and then stabilize in it for 1s (Figure 1C). If participants reached the target within the allotted time, the goal target became green, if they reached the goal target too late it remained red and if they reached it too quickly it turned back to an open circle. The feedback about the timing of reaching movements was provided to encourage consistent movement kinematics but was not used as an exclusion criterion for trials not in the expected time window. Overall, 71.81% of all trials reached the target in the expected time window. However, all trials, whether in the right time window or not, were included in the analysis, as we are interested in the time course of adaptation of participants reaching movements. Moreover, most failed trials reached the target slightly later than the expected time window, often during the initial force-field trials, as participants were facing the perturbation for the first time. Before the task, participants performed a series of 10 trials in the null field to become familiar with the timing requirement.

**Figure 1.**
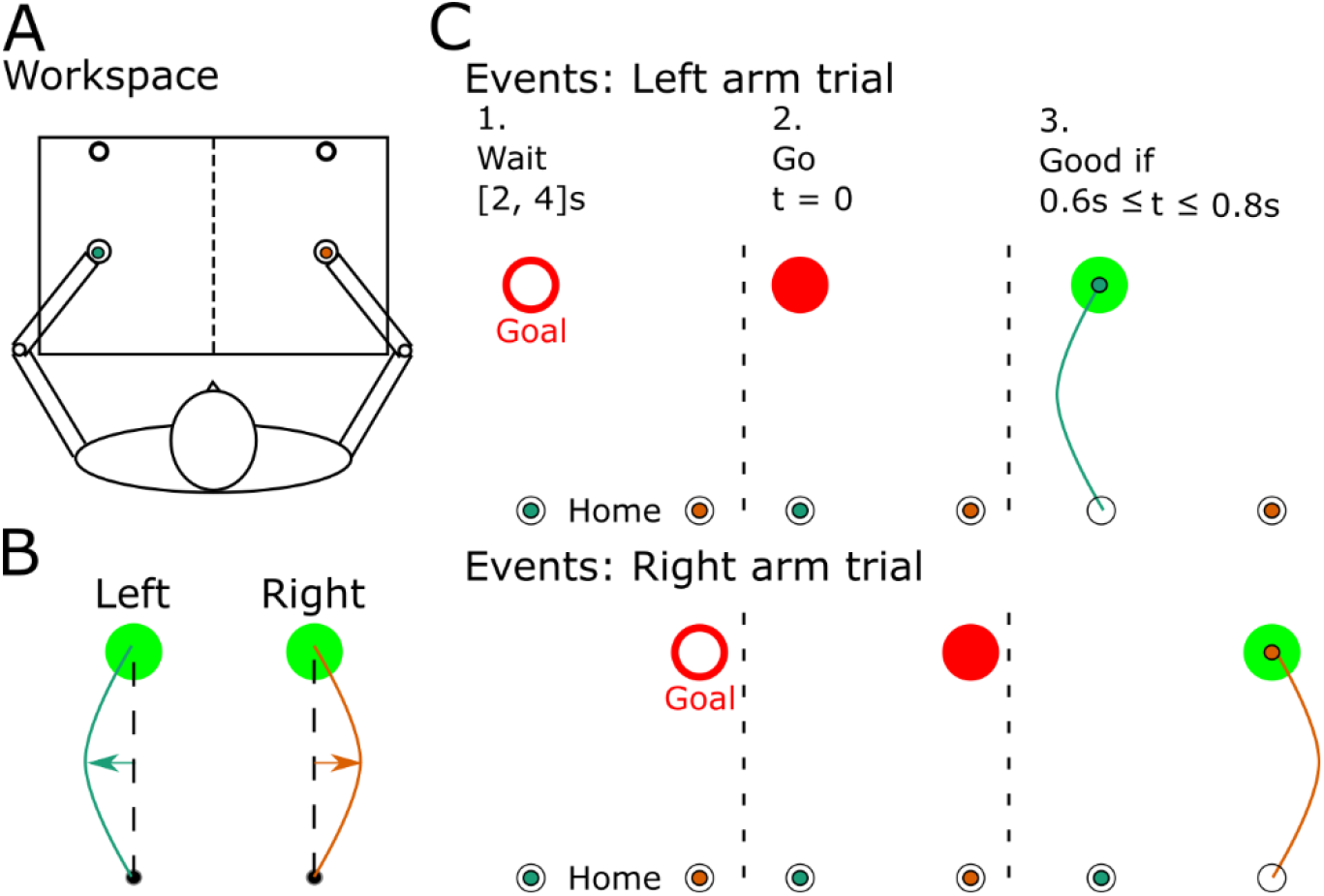
Illustration of the workspace, events during the task, and force field. **A)** Participants were instructed to perform forward reaching movements towards a visual target that was presented in front of either the left or the right arm, with each arm having its own home and goal target. **B)** The force field for experiment 1 had a clockwise direction for the right arm (orange line) and was mirrored for the left arm, hence directing the arm towards the exterior direction. **C)** Events happening during a trial for the left and the right arm. An open goal target was presented for a random period, uniformly distributed between 2 and 4s, before it was filled in. The cue to reach the target was provided by filling the goal target in red. If the participant reached the target in a time comprised between 0.6 and 0.8s, the goal target was filled in a green colour to indicate a good trial and red if the target was reached too slowly.

#### Experiment 1

Experiment 1 aimed at investigating the adaptation of the trajectory and control strategy of the right and left arms in a standard adaptation paradigm (Crevecoeur et al., 2019; Yang et al., 2006). Participants performed force field reaching movements during which the robotic arm applied a curl force field on the participant’s hand. The force field had the form presented in eq. (1):

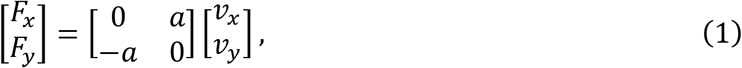

where *F_x_* and *F_y_* represent the x and y force applied on the arm, *v_x_* and *v_y_* represent the x and y velocity of the hand. Parameter *a* was set equal to −13 Nms^-1^ for trials performed with the right hand and 13 Nms^-1^ for the trials performed with the left arm to elicit reactions from the same muscle groups in the two arms (Figure 1B). Null field trials were introduced randomly and used as catch trials to record maximal mirror deviations when the force field was removed unexpectedly. We selected catch trials instead of clamp trials to analyse the feedback corrections during these trials. Participants performed 30 perturbed trials and 5 null field trials with each arm for a total of 70 trials per block. Participants performed 6 blocks of trials for a total of 420 trials. Trials with the left and right arm were randomly presented.

#### Experiment 2

Experiment 2 was designed to analyse rapid changes in feedback control strategies due to unexpected perturbations applied to the dominant and non-dominant arms. Participants performed null field reaching movements with force field trials being randomly interspersed as catch trials. Participants performed the first block of 25 trials with no perturbation with each arm for a total of 50 trials (this block is named BL in the rest of the text). They then performed 6 blocks of 72 trials distributed as follows: 24 null field trials with each arm, 6 force field trials perturbed in the interior direction (clockwise force field for the left arm, and counter-clockwise force field for the right arm) and 6 catch trials perturbed in the external direction (counter-clockwise force field for the left arm and clockwise force field for the right arm). Catch trials occurred with a frequency of 1 catch trial for every 4 null field trials performed with each arm. The force field used during catch trials was of the same form as in Experiment 1 with only the parameter *a* changing from 13 to −13 for clockwise and counter-clockwise force fields. Trials with the left and right arm were also presented in random order.

### Data Analysis

The coordinates of the position of the cursor and the forces measured at the handle were low-pass filtered using a fourth-order dual-pass Butterworth filter with a cutoff frequency of 50Hz. Velocity was obtained from numerical differentiation of position signals. All signals were aligned on movement onset, which was defined as the moment when the cursor position exited the home target.

In the two experiments, surface EMG electrodes (Bagnoli Surface EMG Sensor, Delsys Inc., Natick, MA, USA) were used to record muscle activity during movements. Based on previous studies (Crevecoeur et al., 2019, 2020a), we focused on the Pectoralis Major (PM) and the Posterior Deltoid. Indeed, these muscles have been shown to be strongly recruited to compensate for lateral disturbances and hence should provide valuable information about the strategy employed by participants to counter the perturbation (Crevecoeur et al., 2020b, 2019). EMG signals were sampled at 1KHz and were amplified by a factor of 1000. In both experiments, the reference electrode was attached to the right ankle of the participants. The raw EMG data were band-pass filtered with a 4^th^ order double-pass Butterworth filter with cut-off frequencies set at 20 and 250Hz. EMG data were normalized for each participant to the average activity collected when they maintained postural control at the home target against a constant force of 9N.

To assess the performance of participants during the task, we extracted several key parameters: the maximal deviation of the reaching trajectory (MD), the length of the reaching trajectory, i.e. the path length (PL), and the maximum speed of the reaching movement (MS).

#### Experiment 1

For the force-field trials of experiment 1 we also extracted the maximum force (MF) applied by participants to counter the force field. Moreover, for force field trials we computed the temporal correlation between the lateral commanded force extracted offline based on the forward hand velocity, and the measured force at the handle. This provided us with an index of participants’ motor adaptation to the perturbation (Crevecoeur et al., 2020b). This can be justified as straight movements should exhibit a high correlation since the impact of the commanded force field must be countered by a force equal and opposite. Therefore, changes in the correlation coefficient between the measured and commanded force reflect changes in control, such that increases in the strength of the correlation can be taken as a proxy of adaptation.

In experiment 1, we performed mixed-model analyses on each of these parameters (MD, PL, MS, MF and correlation) with Arm (left and right) and Trial number as fixed effects, and a random intercept for each participant to capture idiosyncrasy. Post-hoc analyses were performed using t-tests with Bonferroni correction for multiple comparisons. To further assess the adaptation of participants across trials, we used a standard, first-order exponential model for learning curves fitted to PL and MD. The exponential fit was defined as follows:

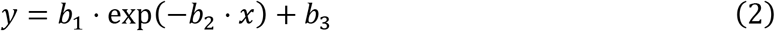

Where *y* represents the dependent variable (either PL or MD), *x* is the trial number and *b*_1_, *b*_2_ and *b*_3_ are the parameters of the fit. Parameter *b*_2_ [1/trial] represents the learning rate of participants as it defines the number of trials participants needed to attain stable performance. The parameters of the fit were compared across the left and right arms by performing a bootstrap analysis. The participants were resampled with replacement, and the parameters of the exponential fit were calculated for each bootstrapped population. This allowed us to derive a distribution of parameters across the 1000 bootstrapped samples and compare these distributions across limbs. Finally, to determine whether participants used co-contraction as a strategy to counter the force field, we computed the average EMG level in a window of 100ms before movement onset for the 10 first and the 10 last trials. As EMG data has more variability than kinematic data, we chose 10 trials to have a good measure of the average EMG behavior. In this experiment, three of the twelve participants were excluded from the EMG analysis due to problems with the EMG recordings. Paired t-tests were used as Post-hoc analysis and in the case of non-significant results we computed the Bayes Factors (BFs). BF stands for the ratio between the likelihood of H1 (there being a significant difference between the two populations) over the likelihood of H0 (there being no significant difference between the two populations). A BF between 0.33 and 1 provide small evidence of no difference, a BF between 0.1 and 0.33 provides substantial evidence, and a BF < 0.1 indicates strong evidence (Keysers et al., 2020).

#### Experiment 2

In experiment 2, trials before the introduction of the force field were considered baseline trials. As for experiment 1, we used a linear mixed model analyses on each of the extracted parameters (MD, PL, and MS) with Arm (left and right) and force field presence (FF) as fixed effects, and a random intercept per participant. We also extracted MS on trials immediately following a catch trial and performed a linear mixed model analysis with Arm and Trial number as fixed predictors. Finally, for the catch trials, we performed a paired-wise t-test with the arm as the test variable. All post-hoc analyses were performed using Bonferroni adjusted t-tests.

Finally, to determine whether participants used co-contraction, we computed the average EMG level in a window of 100ms before movement onset for the 10 first and 10 last null field trials of experiment 2. We performed a repeated measure ANOVA with arm and position (Early versus late adaptation) as within factor parameters. Paired t-tests were used as Post-hoc analysis and in the case of non-significant results we computed the BFs.

## Results

### Experiment 1

In experiment 1 participants were exposed to force field perturbations, with the left arm exposed to a counter-clockwise force field and the right arm exposed to a clockwise force field. The performance of participants improved during each block, resulting in trajectories with significant lateral deviations in the first trials that became more rectilinear as trials progressed (Figure 2A). We extracted PL, MD, and MS of the hand trajectories to measure changes in reaching movements across trials.

**Figure 2.**
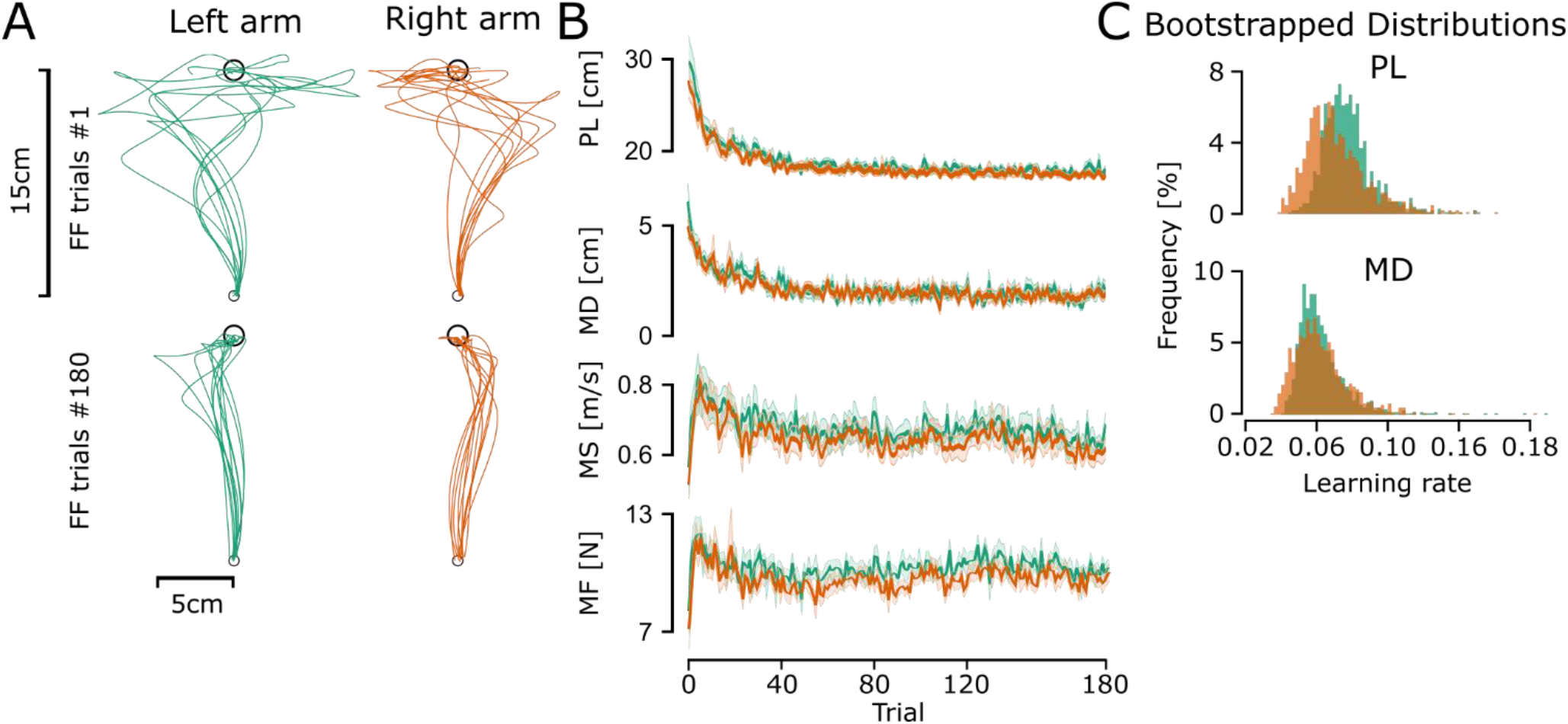
First and last trajectories for the force field perturbed trials, the evolution of parameters across all trials, and histograms of the learning rate of path length and maximal deviation. **A)** First and last force field trials for the left arm (green line) and the right arm (orange line) of participants. **B)** Mean and standard error of the mean of Path length (PL), maximal deviation (MD), maximum speed (MS) and maximum force (MF) across perturbed trials of experiment 1 for the right arm (orange line) and the left arm (green line). **C)** Bootstrapping results of the learning rate (parameter *b*_2_ in eq. (2)) of the path length and maximal deviation for the left and right arm of all force field trials and all participants

PL and MD decreased as the number of trials increased (Figure 2B), thereby highlighting that participants adapted their reaching movements in the presence of the force field. A linear mixed model analysis showed that PL (Marginal R^2^ = 0.45 and conditional R^2^ = 0.557) presented a significant effect of arm (estimate = −0.007, F = 28.16, p < 0.001), a significant effect of trial number (estimate = −0.00025, F = 1305.1, p < 0.001) and a significant interaction effect (estimate = 0.00003, F = 5.17, p = 0.023). Similarly, MD (Marginal R^2^ = 0.27 and conditional R^2^ = 0.43) presented a significant effect of arm (estimate = −0.0025, F = 30.10, p < 0.001) and trial number (estimate = −0.000075, F = 6.3876, p = 0.01) but no significant interaction (estimate = 0.00001, F = 1.79, p = 0.18). The significant effect of trial number on both PL and MD indicates that participants performed straighter movements as trials progressed (Figure 2A). The significant effect of arm indicates that the left arm had a longer PL (estimate = 7mm) and was deviated more (estimate = 1.5mm) across all trials, while the significant interaction indicates differences in learning rates for PL between the two arms.

MS presented a different evolution across trials, increasing during the first trials, reaching a maximum rapidly, and then decreasing for the remaining trials (Figure 2B). A linear mixed model (Marginal R^2^ = 0.099 and conditional R^2^ = 0.36) showed a significant effect of arm (estimate = −0.032, F = 23.4535, p < 0.001), trial number (estimate = −0.0004, F = 152.16, p < 0.001) and no significant interaction (estimate = 0.000008, F = 0.0165, p = 0.89). This indicates the arms performed the reaching movements at different speeds, with the left arm having a higher MS across all trials (estimate = 0.032m/s). Moreover, both arms displayed a significant but similar change across trials (Figure 2B).

The maximum force followed a similar evolution to MS (Figure 2B), linear mixed model analysis (Marginal R^2^ = 0.018 and conditional R^2^ = 0.032) of the maximum force applied by each arm to counter the perturbation during force field trials showed a significant effect of Arm (F = 28.3721, p < 0.001), with the left arm applying more force and no significant effect of trial number (F = 0.27, p = 0.6) and no significant interaction (F = 0.45, p = 0.5). Indeed, the left arm produced a higher average peak force to counter the perturbation (10.4066 +-1.91N) than the right arm (9.8392 +-2.02N).

Post-hoc analysis of the difference between the left and right arm of the first 5 and last 5 trials highlights that there was a significant difference in PL in the first 5 trial (t = 2.3293, df = 57, p = 0.02, d = 0.31) with the left arm having a larger PL than the right arm, and no significant difference for the last 5 trials (t = 1.69, df = 59, p = 0.09, d = 0.22). For MD, we found no significant difference in the first 5 trials (t = 1.863, df = 57, p = 0.067, d = 0.24) or the last 5 trial (t = 1.67, df = 59, p = 0.09, d = 0.22). This indicates that a difference exists in the first perturbed reaching movements of the two arms, with the right arm being less perturbed by the force-field and presenting a straighter trajectory (Figure 2A). The difference disappeared in the late trials where participants displayed similar deviations across the arms (Figure 2A). A paired t-test comparing MS of the first 5 trials showed a non significant difference between the two arms (t = 1.87, df = 57, p = 0.065, d = 0.25), whereas a paired t-test of the last 5 trials showed a significant difference between the right and left arm (t = 3.077, df = 59, p = 0.003, d = 0.4, mean of the differences = 0.03 m·s^-1^) confirming that the left arm had a greater maximum speed during the last trials (Figure 2B). This is important because it already highlights different control strategies: indeed, as the force was proportional to velocity, the perturbation force was larger to the left arm, although we have seen above that there were no differences between the two arms in PL and MD in the last five trials.

We performed an exponential fit on PL and MD to determine the rate of learning (*b*_2_ in eq. (2)) of each arm. We performed this exponential fit on all participants together and we performed a bootstrap analysis with resampling of participants with repetition to determine the distribution of the parameters of the fit (Figure 2C). A paired t-test comparing the two arms showed a significant difference in the learning rate for both PL (t = 11.129, df = 999, p < 0.001, d = 3.41) and MD (t = 2.5015, df = 999, p < 0.001, d = 0.45), with the left arm learning at a faster rate (Figure 2C). More precisely, we observe differences in learning rate that suggest faster adaptation in the left arm, however, the difference was small.

These results highlight that both arms had similar performances during the last trials, with similar maximal deviations and a slightly larger PL for the non-dominant arm. However, the non-dominant arm did perform the task at a higher speed and used a larger force to counter the perturbation during the task.

To further characterize how participants adjusted their reaching movement to the force field specifically, we analyzed the correlation between the force applied by the participants on the robot handle (F_x_ in Figure 3A) and the force applied by the force field (proportional to Vy in Figure 3A). We can observe that during the first trial, F_x_ was not well correlated with Vy for both the left and right arms (Figure 3A), the correlation improved across trials and both signals became more closely correlated during the late adaptation phase (Figure 3A). The correlation between F_x_ and Vy improved across trials (Figure 3B) with the correlation of the left arm going from 0.5689 ± 0.204 in the first trial to 0.84 ± 0.0814 in the last trial, and from 0.4576 ± 0.1968 to 0.885 ± 0.033 for the right arm. A linear mixed model analysis (Marginal R^2^ = 0.41 and conditional R^2^ = 0.58) showed a significant effect of arm (estimate = 0.035, F = 60.75, p < 0.001) and trial number (estimate = 0.00064, F = 1003, p < 0.001) and no interaction (estimate = 0.00007, F = 1.803, p = 0.0715). On the one hand, the significant effect of trial number highlighted the increase in correlation across trials, which indicates that both arms adapted to the presence of the force field. On the other hand, the significant effect of the arm indicated different levels of correlation across trials. We compared the average correlation of the last 20 trials of the left arm and the right arm for each participant to analyze whether participants showed greater improvement with the dominant or the non-dominant arm. A paired t-test showed a significant effect of Arm (t = −10.55, df = 251, p < 0.001, d = −0.66, Figure 3B and C) on the correlation for the last 20 trials. The dominant arm presented a higher average correlation across the last 20 trials for eleven of the twelve participants, which indicated that participants adapted to the perturbation more with their dominant arm (Figure 3C). Furthermore, we computed the Pearson’s correlation coefficient to determine whether participants presenting a good adaptation with one arm also presented a good adaptation with the other arm. This correlation highlighted that participants with a good adaptation on one arm were also good with the other arm (r = 0.7464, p = 0.005, Figure Catch trials where no perturbation was applied were randomly interleaved during experiment 1 to observe the after-effects of the adaptation of control strategies of the two arms. The absence of a force field caused participants to deviate in the direction opposite to the force field (Figure 4A). All three variables (MD, PL and MS) showed a difference across arms (Figure 4B-D). A paired-wise t-test showed a significant difference for MD (t = 6.6146, df = 354, p < 0.001, d = 0.35), PL (t = 4.0308, df = 354, p< 0.001, d = 0.21) and MS (t = 3.7145, df = 354, p < 0.001, d = 0.2). Qualitatively similar results were observed when the statistical tests were performed on subject average. This shows that the dominant arm was less impacted by the introduction of catch trials, while the non-dominant arm used a higher forward speed than the right arm.

**Figure 3.**
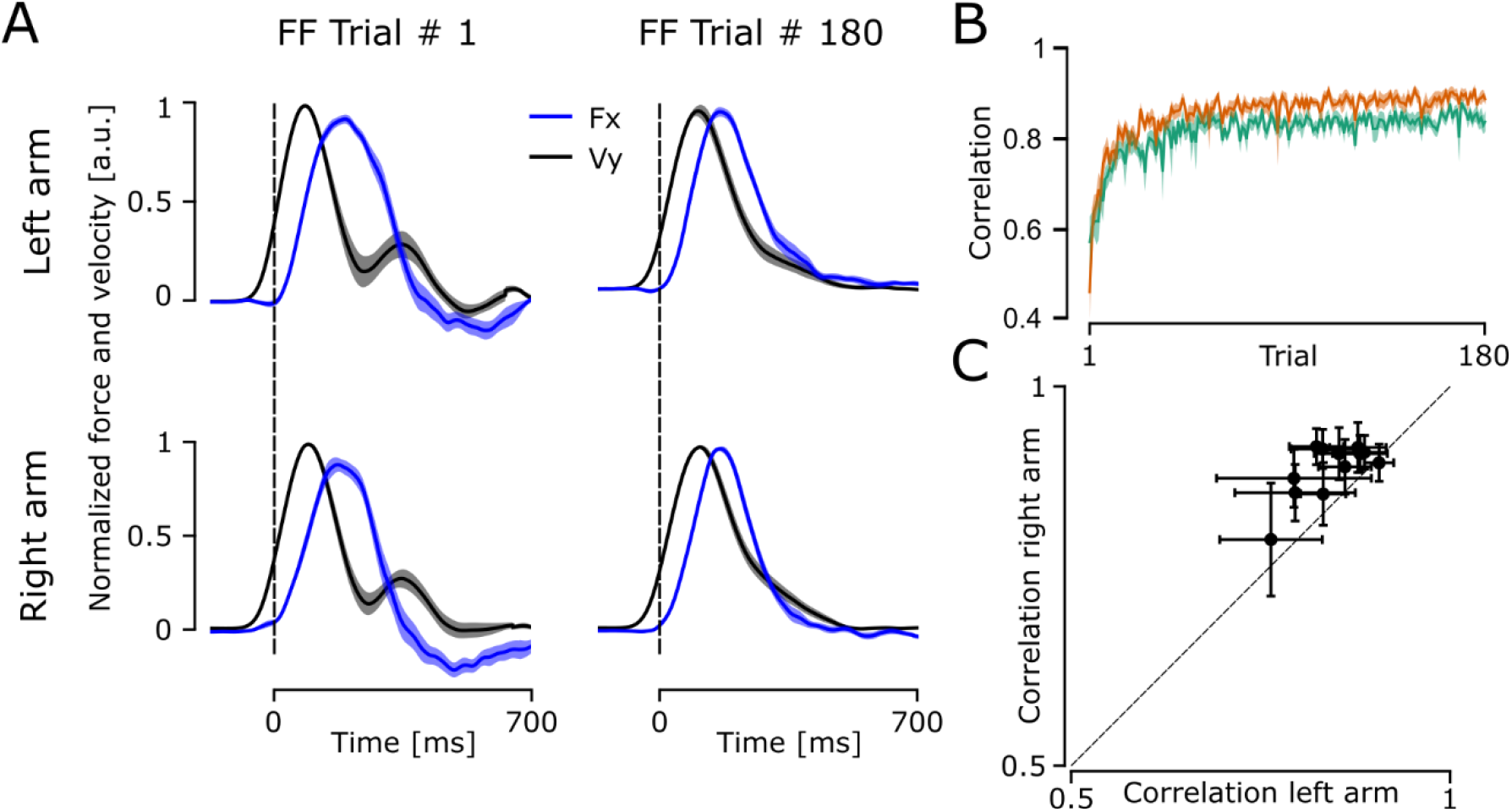
Correlation between the force exerted by participants on the handle of the robotic arm and the force exerted by the robot on the participants’ arms. **A)** The mean and standard error of the mean of the normalized x force applied by the participant on the handle (blue line) and the normalized y velocity of the reaching movement (black line) for the first and last perturbed trials. The force and velocity were normalized to their peak value and averaged across all participants. **B)** Mean and standard error of the mean of the correlation between force and velocity across all perturbed trials for the left arm (green) and the right arm (orange). **C)** Correlation average for the final 20 trials for the left and the right arm of each participant (black dots), horizontal and vertical error bars represent the standard deviations of the correlation of the left arm and the right arm respectively.

**Figure 4.**
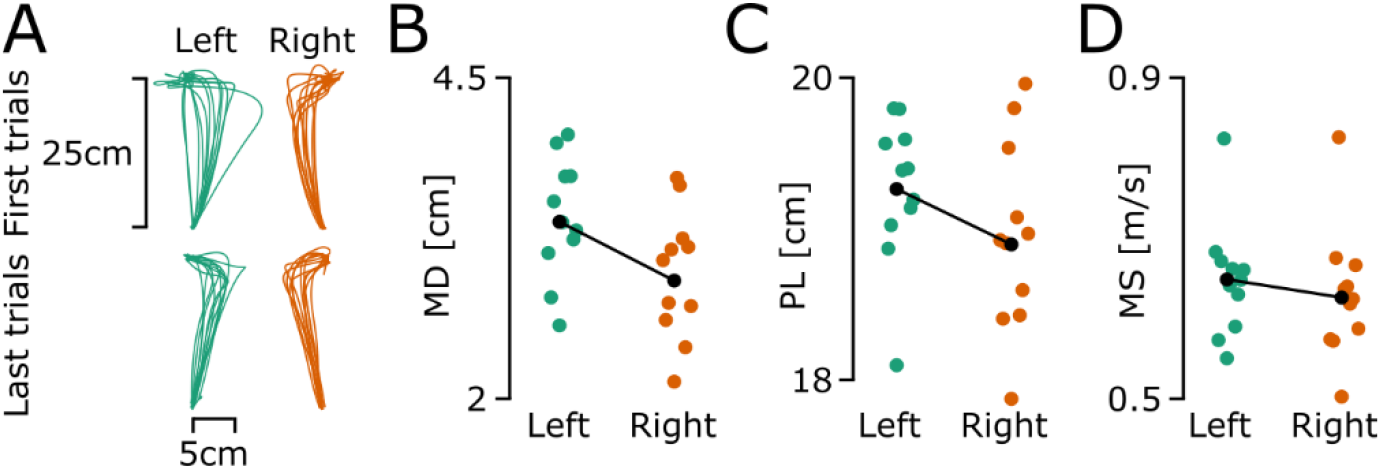
Trajectories of first and last catch trials of experiment 1 for all participants and mean maximal deviation, path length, and maximum speed of participants across all catch trials of experiment 1. The left arm is presented using a green colour across all panels and the right arm is presented using orange colour. In panels B to D The global average for each arm is presented as a black dot. **A)** Trajectories of the first and last catch trials for all participants. **B**) Maximal deviation across all catch trials. **C)** Path length across all catch trials. **D)** Maximum speed across all catch trials.

### Experiment 2

Experiment 2 was designed to probe feedback responses to unexpected force fields and investigate possible differences in participants’ ability to adapt the feedback response of an ongoing movement (Crevecoeur et al., 2020a). They performed null field reaching movements to a target presented 15cm away from the starting positions. After twenty-five null field trials with each arm (Baseline, BL), force field trials, in either the clockwise or counter-clockwise direction, were introduced on one out of five trials. The force field trials and null field trials were interleaved in random order. During baseline trials participants’ performance was stable. Once force field trials were introduced MD and PL showed small increases while MS increased more significantly for both arms in null field trials (Fig. Figure 5A). A change in MS has been linked to a change in the control strategy used to perform reaching movements (Crevecoeur et al., 2019). Therefore, to examine whether participants used a more robust control strategy following the introduction of force field trials, we performed a linear mixed model analysis on MS with Arm and force field (FF) as within-trials parameters. The linear mixed model had a marginal R^2^ = 0.07 and a conditional R^2^ = 0.3. We found a significant effect of FF (F = 404.276, p < 0.001), no significant effect of arm (F = 0.0737, p = 0.786) and no interaction (F = 0.3574, p = 0.55). A post hoc analysis showed no significant difference in MS between the two arms, before and after the introduction of force field trials (before force field: p = 0.786; after force field: p = 0.433). This result shows that participants adapted their control gains without a clear difference between the two arms.

To assess the influence of force field trials on the control strategy in more detail, we measured the change in MS on the trials immediately following a force field trial (Figure 5B). MS increased after a force field trial and slowly diminished in the following trials without ever coming back to the levels observed during the baseline trials. A linear mixed model analysis (Marginal R^2^ = 0.05 and a conditional R^2^ = 0.22) showed a significant effect of Arm (F = 6.23, p = 0.0126) and Trial number (F = 62.17, p < 0.001) and no interaction (F = 0.4217, p = 0.52). This suggests that MS changed across trials and that both arms showed different maximum speeds across baseline and trials following force field trials. A post-hoc analysis showed a significant difference in MS between baseline trials and trials following a force field trial for both the right arm (FF+1 – BL = 0.07, CI = [0.05; 0.1] m/s, p < 0.001) and the left arm (FF +1 – BL = 0.09, CI = [0.06; 0.12], p < 0.001) confirming the increase in MS after a force field trial. Further analysis showed a significant difference between the right and left arm for trial fft+1 (p = 0.008) and no significant difference for other trials (p > 0.139). Concerning the PL, a linear mixed model analysis (Marginal R^2^ = 0.018 and a conditional R^2^ = 0.18) showed a significant effect of Arm (F = 30.9499, p < 0.001), no significant effect of Trial number (F = 0.0004, p = 0.98) and no significant interaction (F = 0.16, p = 0.6867). This highlights that the non-dominant had a trajectory that was slightly more curved than the dominant arm, but the difference was less than 1mm in maximal deviation (see Figure 5B).

**Figure 5.**
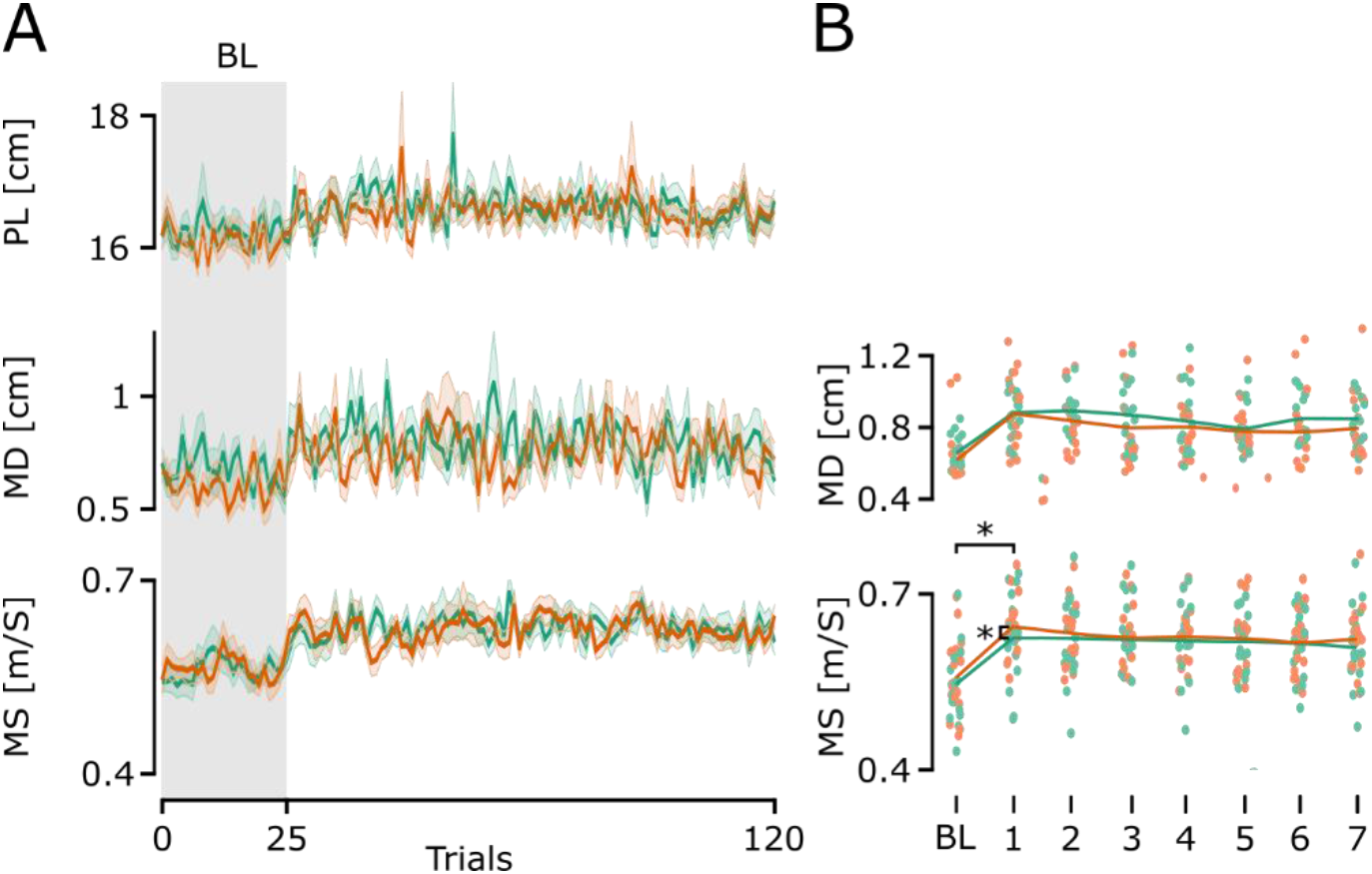
Evolution of extracted parameters in the null field trials in experiment 2. **A)** Mean and standard error of the mean of the path length, maximal deviation, and maximal speed of the reaching trajectories for the left arm (green line) and the right arm (orange line) across all trials. The grey zone indicates baseline trials without force field trials. **B)** Mean MS and MD for baseline trials and trials following force field trials for each participant. The x-axis shows the index of the trials after a force field trial.

Looking at the adaptation to the randomly interleaved force field trials, participants showed an adaptation across trials as shown by an increase in the correlation between the force applied by the robot and the force produced by the participant (Figure 6A and B). For force field trials with the same direction as in experiment 1 (Figure 6A), participants showed a greater correlation with the right arm when compared with the left arm (Figure 6C and E), whereas this difference was not present during force fields in the other direction (Figure 6D and F). A linear mixed model analysis of the inward force fields (marginal R^2^ = 0.024 and a conditional R^2^ = 0.27) shows a significant effect of the trial (F = 15.859, p < 0.001), no significant effect of the arm (F = 1.54, p = 0.215), and no significant interaction (F = 0.18, p = 0.66). This confirms the feedback adaptation across trials and the lack of difference between the arms (Figure 6F). For the outward force fields, the linear mixed model analysis (marginal R^2^ = 0.098 and a conditional R^2^ = 0.38) showed a significant effect of the arm (F = 23.495, p < 0.001) and trial (F = 67.124, p < 0.001) and no interaction (F = 1.423, p = 0.23) which confirms the better adaptation of the dominant arm when compared to the non-dominant arm (Figure 6E). Altogether, these results suggest that the right arm shows a greater adaptation of its behaviour to the presence of a force field.

**Figure 6.**
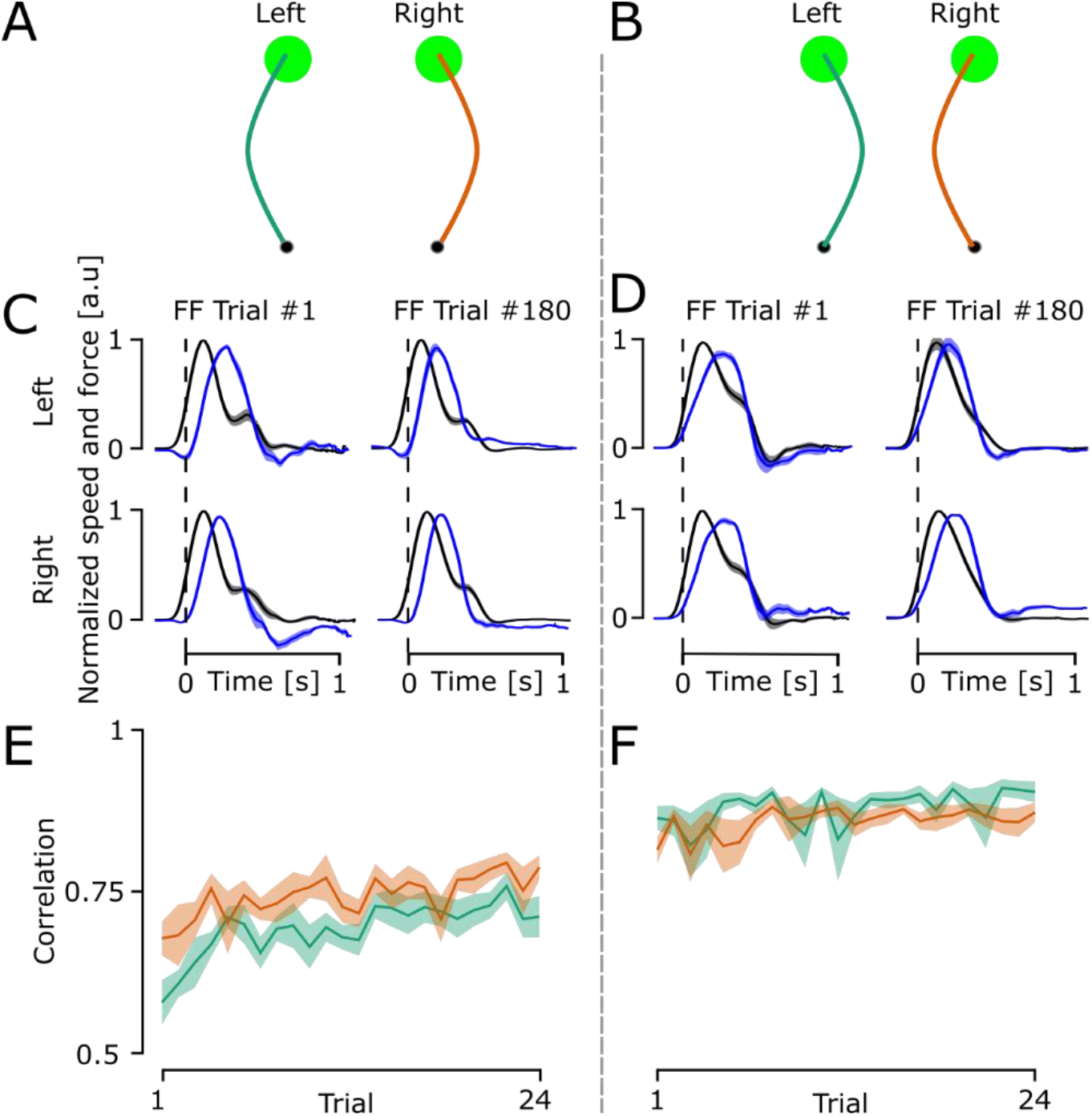
Outputs of the force field trials of experiment 2. A) Representative traces for outward force field trials for the left arm (green line) and the right arm (orange line). The hand paths shown in the figure are for demonstration purposes and do not represent real data. B) Representative traces for inward force field trials for the left arm (green line) and the right arm (orange line). C) The mean and standard error of the mean of the normalized x force applied by the participant on the handle (blue line) and the normalized y velocity (black line) of the outward force field trials reaching movements for the first and last perturbed trials. The force and velocity were normalized to their peak value and averaged across all participants. D) The mean and standard error of the mean of the normalized x force applied by the participant on the handle (blue line) and the normalized y velocity (black line) of the inward force field trials reaching movements for the first and last perturbed trials. E) Mean and standard error of the mean of the correlation between force and velocity across all perturbed trials for the left arm (green) and the right arm (orange) for outward force field trials. F) Mean and standard error of the mean of the correlation between force and velocity across all perturbed trials for the left arm (green) and the right arm (orange) for inward force field trials.

### Co-contraction

We extracted the mean EMG signal for PD and PM during a period 100ms before movement onset to examine whether participants used muscle co-contraction to modulate the limb intrinsic properties and counter the force field (Figure 7). This time window was selected as impedance control is assumed to be effective if active prior to movement. Indeed, if measured during movement, changes in EMG are confounded with feedback control (Franklin et al., 2008).

**Figure 7.**
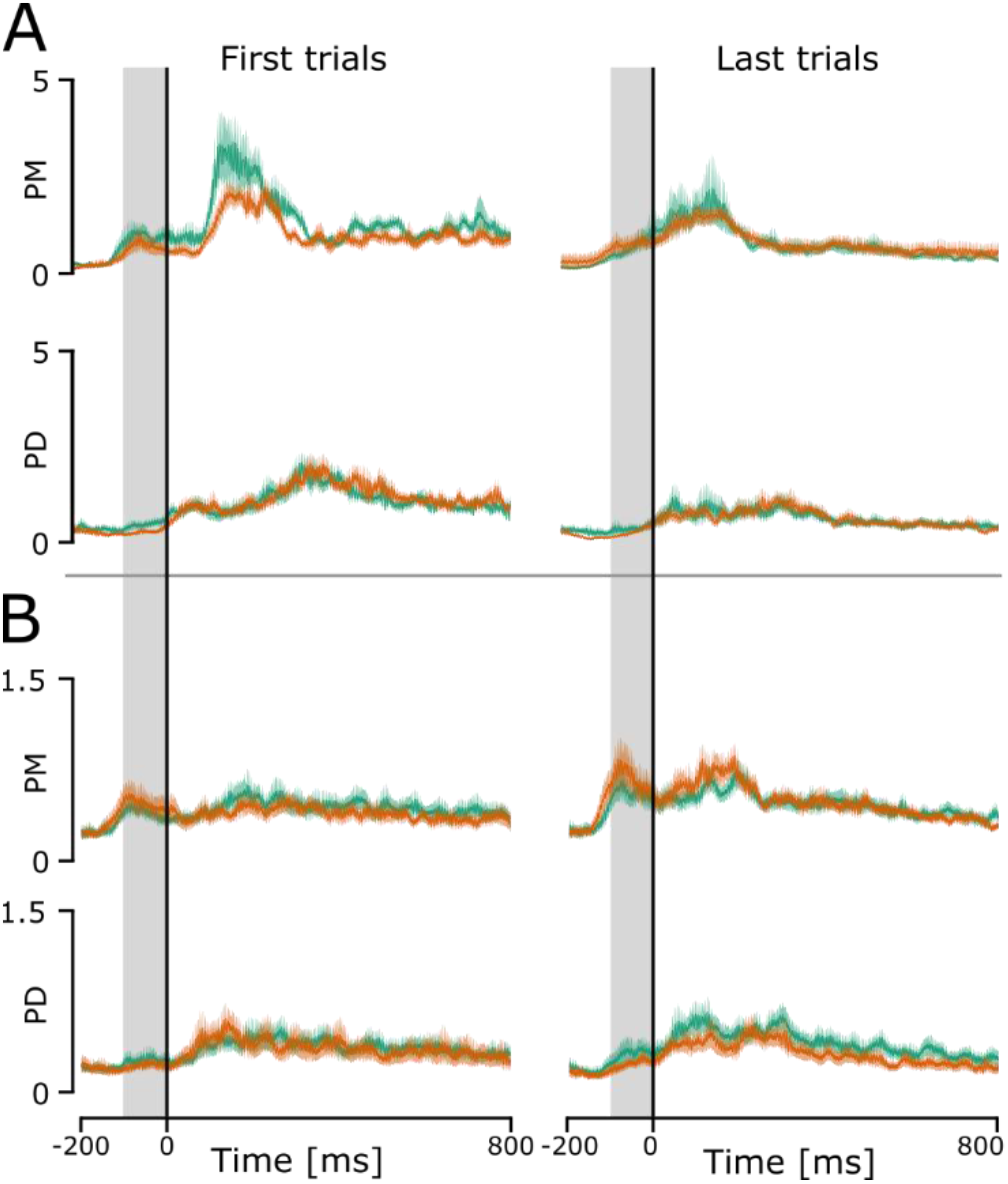
Mean and standard error of the mean of normalized EMG of the posterior deltoid and the pectoralis major averaged across all participants for the right arm (orange line) and the left arm (green line). The mean EMG for both muscle groups was computed in a 100ms window before movement onset (the grey region in the plot). A) Mean and standard error of the mean of the normalized EMG of the 10 first and 10 last force field trials in experiment 1. B) Mean and standard error of the mean of the normalized EMG of the 10 first baseline trials and 10 last null field trials after the introduction of force field trials in experiment 2.

In experiment 1, we extracted these values for the first 10 and last 10 force field trials (Figure 7A). Three of the twelve participants were excluded from the analysis due to EMG electrodes disconnecting during the experiment. A linear mixed model analysis on PD (marginal R^2^ = 0.052 and a conditional R^2^ = 0.45) showed no significant effect of trial (F = 0.2235, df = 30, p = 0.64), no significant effect of arm (F = 3.8017, df = 30, p = 0.06) and no interaction (F = 0.132, df = 30, p = 0.72). A post hoc analysis showed no difference between left and right arm for the first 10 trials (95% CI = [−0.301, 1.35], p =0.1846, BF = 0.5) and no significant difference between the two arms for the last 10 trials (95% CI = [−1.15, 3.35], p = 0.2977, bf = 0.5). Similarly, a linear mixed model analysis on PM (marginal R^2^ = 0.067 and a conditional R^2^ = 0.495) showed no significant effect of trial (F = 1.934, df = 30, p = 0.17), no significant effect of arm (F = 2.33, df = 30, p = 0.14) and no interaction (F = 1.454, df = 30, p = 0.2373). A post hoc analysis showed no difference between left and right arm for the first 10 trials (95% CI = [−0.09, 1.01], p =0.09, BF = 1.1) and no significant difference between the two arms for the last 10 trials (95% CI = [−0.606, 0.36], p = 0.5894, bf = 0.35). These results indicate that cocontraction was not used as a strategy to improve performance when facing the force field.

In experiment 2, we computed these same values for the first ten baseline trials and the last ten null field trials (Figure 7B). A linear mixed model analysis for PD (marginal R^2^ = 0.032 and a conditional R^2^ = 0.67) showed a significant effect of arm (F = 4.22, df = 48, p = 0.045), no significant effect of force field presence (F = 2.23, df = 48, p = 0.1419) and no significant interaction (F = 0.5887, df =48, p = 0.55). A post hoc analysis showed no difference between left and right arm for the first 10 trials (95% CI = [−0.04, 0.13], p =0.311, BF = 0.41) and no significant difference between the two arms for the last 10 trials (95% CI = [−0.01, 0.17], p = 0.066, bf = 0.63). For PM, the linear mixed model (marginal R^2^ = 0.06 and a conditional R^2^ = 0.57) showed a significant effect of force field presence (F = 1.73, df = 48, p = 0.008), no significant effect of arm (F = 1.73, df = 48, p = 0.195) and no interaction effect (F = 0.091, df = 48, p = 0.764). A post hoc analysis showed a significant increase between the first ten baseline trials and last ten null-field trials for the right arm (95% CI = [−0.39, −0.02], p = 0.034, d = −0.43, bf = 0.31) and no significant difference for the left arm (95% CI = [−0.35, 0.02], p = 0.086, bf = 0.36). Altogether, these results suggest that the changes in performance in both experiment 1 and experiment 2 were not related to co-contraction modulating the limb intrinsic properties.

## Discussion

We explored control and adaptation strategies to investigate differences linked to hand dominance. We found that participants adapted their behaviour in the presence of a force field (experiment 1) with very similar patterns of adaptation across the two arms. We observed that the non-dominant arm presented a greater forward speed, a greater reactive force that was a direct consequence of the force field definition, similar deviation, and reduced correlation when compared with the dominant arm. We also showed that when confronted with randomly interleaved force field trials (experiment 2) participants increased the speed of their reaching movement even during trials where no force field was present. Moreover, in experiment 1 we observed that participants showed a better adaptation with their dominant arm with a good correlation between the adaptation of the two arms (Figure 3C). We did not observe any main change in co-contraction that could explain the changes in behaviour observed across early and late phases of exposure to disturbances in both experiments, although trial-by-trial modulation of co-contraction could have been evoked transiently by the force field of Experiment 2.

Handedness is often understood as a specialization of the role of each arm. It has been suggested that the dominant arm relies more heavily on “predictive” control, using internal models of limb dynamics more efficiently (Bagesteiro and Sainburg, 2002) while the non-dominant arm relies on impedance control mechanisms (Bagesteiro and Sainburg, 2003; Jayasinghe et al., 2022; Mutha et al., 2013; Woytowicz et al., 2018; Yadav and Sainburg, 2014b). This hypothesis suggested that the dominant arm is more specialized in the control of movements whereas the non-dominant arm is more specialized in the control of arm postures. The specialization of each arm has also been linked to the specialization of the left and right hemisphere of the brain, with the hemisphere contralateral to the dominant arm being specialized in predictive control of limb dynamics and the other hemisphere being specialized in controlling the limb impedance. Indeed, papers studying deafferented patients showed lateralized differences in the effect of the lesion in different aspects of control (Schaefer et al., 2012, 2009, 2007). They observed that damage to the left hemisphere led to deficits in trajectory control, whereas as damage to the right hemisphere led to deficits in final position control. More recently, studies on somatosensory deafferentation also revealed lateralized roles of proprioceptive feedback (Jayasinghe et al., 2020, 2021), with the right arm failing to stabilize at the end-point of the reaching movement and the left arm showing poor corrections of the trajectory. However, other results have shown that the two arms can develop feedforward adaptation equally well in an experiment in which participants performed fast straight reaching movements, with similar force fields to our experiments (Reuter et al., 2016). We tested the specialization hypothesis with comparisons of the two arms of right-handed participants in two experimental paradigms where the trials with each arm were randomly interleaved. This approach allowed us to focus on individual differences within participants and enabled a direct comparison of the adaptation mechanisms used by the two arms. Our data do not provide support for this previous model. Indeed, concerning predictive aspects, we found that the two arms adapted very well in parallel with comparable movement parameters and learning rates, similar to Reuter et al. (2016). Moreover, both arms displayed similar reductions in maximal deviation and PL (Figure 2). In Experiment 1, differences were highlighted in the speed of the movement, the force and the correlation between the perturbing force applied by the robotic arm and the reactive force produced by the participant, which suggested a better adaptation to the force field of the dominant arm (Figure 3).

During the initial trials of the task, participants have no knowledge of the force field that will be applied to their arm. We consider the problem of reaching without an accurate model of the perturbation in the context of robust control (*H*_∞_ control). Robust control and linear quadratic gaussian (LQG) control are mathematically very similar but have one significant difference. On the one hand, LQG assumes that disturbances in the system can be modeled by white Gaussian noise with known covariance matrices (Todorov and Jordan, 2002). On the other hand, robust control does not make any assumption on the exogeneous perturbation signal, leading to a control solution that aims to minimize the impact of a ‘worst-case’ perturbation on the system. In reaching movements this results in an increase in control gains that produces larger speed and more vigorous responses to external loads (Crevecoeur et al., 2019). Of course, a controller can be placed in a spectrum between optimal control, which assumes a perfect model of the perturbation, and robust control, which assumes no model of the systematic force disturbances.

It has been shown that a modulation of reaching speed when confronted to a force field is linked to the use of a more robust control strategy (Crevecoeur et al., 2019). In our case, we observed an increase in movement speed across trials in both experiments 1 and 2. In experiment 1, the robust strategy is highlighted by considering that the movement speed and the maximum force in the nondominant arm were larger, yet the deviation was comparable. Because the force field was proportional to velocity, this means that they used larger feedback gains without relying on a more accurate model since the continuous correlations between commanded and measured forces were reduced in the non-dominant arm. Thus, their strategy consisted of a stronger disturbance rejection that did not rely on an accurate model of the force field, which corresponds to robust control. In experiment 2, we also observed that MS of null field reaching movements for both arms increased once force field trials were introduced (Figure 5B) with no difference between the two arms. This suggests that both arms relied on a comparable modulation of the robustness of control in the random context of experiment 2.

Interestingly, in experiment 1 we also observed that the non-dominant arm was more impacted by the catch trials as shown by a greater PL, PL, and MS (Figure 4). This could result from the greater speed and force of the reaching movement of the non-dominant arm during force field trials as well as the less accurate representation of the force field (smaller correlation in Figure 3B). Indeed, when performing reaching movements, participants are likely expecting a higher perturbation force for the non-dominant arm when compared to the dominant arm, leading to a greater perturbation when the force field was unexpectedly removed.

Robust control differs from impedance control on two aspects. First, robust control assumes errors (or uncertainties) in the accuracy of the representation of the perturbation leading to an increased speed and vigorous response to perturbations (Crevecoeur et al., 2019). Impedance control assumes that an increase in the contraction of agonist and antagonist muscles will lead to a greater stabilization of the joint (Hogan, 1984). Therefore, we expect the robust controller to display an increase in control gains, resulting in faster movements toward the target and more vigorous responses to perturbations. Our data shows that the neural controller was more robust in the sense that the increase in feedback responses reduced the impact of the perturbations with less adaptation. All of this without assuming that such a response is obtained through the activation of agonist and antagonist muscles. In experiment 1, the occurrence of force field disturbances evoked both faster movements and more vigorous responses to perturbations. With the non-dominant arm showing both higher movement speed and higher force to counter the perturbation when compared with the dominant arm, suggesting the use of a more robust controller for the non-dominant arm. In experiment 2, we observed an increase in MS in both arms after force-field trials were introduced, suggesting the use of a more robust controller. Hence, our results suggest a robust control strategy, which can be dissociated from automatic stiffening of the limb through impedance control (Etienne Burdet et al., 2001; Hogan, 1984). Indeed, we didn’t observe a particular increase in muscle co-contraction and such an increase in co-contraction could have only moderately altered the intrinsic properties of muscles (Crevecoeur and Scott, 2014b).

What our interpretation adds to the field can be understood in terms of the quality of internal representations used to perform reaching movements. First, we emphasize feedback control models because a clear difference arose in the correlation which includes online compensation throughout the whole movement (Figure 3). From a computational perspective, optimal control models have often assumed that movement disturbances followed Gaussian distributions (Todorov and Jordan, 2002), without explicitly formulating the problem of control with unmodelled disturbances. Such unmodelled disturbances may involve model errors, or errors due to novel environments, in which case, feedback control can either compensate for this disturbance without knowledge, which is the purpose of a robust controller, or learn about the novel dynamics and adjust control accordingly (i.e. adaptation). The ability to derive novel optimal control laws following adaptation thus depends on the ability to acquire a novel and accurate representation of the dynamics of a novel force field. We showed that both arms were able to do so, with a small advantage for the dominant arm. Moreover, in this framework, the fact that the non-dominant arm made greater use of a robust policy may reflect that the novel internal model on this side was slightly less accurate.

Interestingly, participants who showed a greater correlation with the dominant arm also showed a greater correlation with the non-dominant arm (Figure 3B). This is in line with results from (Maurus et al., 2021) who argued that the internal model of limb dynamics was similar across arms. Moreover, Maurus et al. showed that sensory feedback has a similar role for the dominant and the non-dominant arm when countering random disturbances with the arms during postural control. In our experiments, we aimed at minimizing the potential transfer of knowledge between one arm and the other by randomizing the order of the arm performing the reaching movement during our task. Therefore, we think that it is unlikely that transfer fully accounts for the relationship between correlation levels across participants. However, it could still account for some part of the observed effect as it has been shown that interlimb transfer is asymmetrical, with most of the learning transfer going from the dominant to the non-dominant arm (Galea et al., 2007; Wang and Sainburg, 2004), limited in magnitude (Mostafa et al., 2014; Taylor et al., 2011), and its magnitude is not affected by the training schedule (Joiner et al., 2013). Our results indicate that both arms of all participants share a common mechanism of adaptation.

In experiment 2 we also observed difference between the two force field directions, with participants showing a greater adaptation to the perturbation for both arms for the inward oriented force field than for the outward oriented force fields (Figure 6). These differences could arise from directional preferences of movements and on the muscles that are active in countering the force field perturbations. Indeed, biomechanical considerations have been shown to impact the choice of the arm used to perform reach movements (Bryden and Roy, 2006), and of preferential direction of force generation during bimanual tasks (Córdova Bulens et al., 2018). Such preferential directions could lead to the differences in the response to the perturbation observed in experiment 2, with participants having a greater ease to react to perturbations driving the arm inwards.

It has to be noted that participants were self-assessed right-handed and therefore could present different degrees in laterality. Indeed, previous studies have shown that the degree of laterality can impact the transfer of learning across the two arms (Lefumat et al., 2015). However, the use of a random trial schedule in our study should minimize the transfer of learning. Moreover, only right-handed participants performed the experiment, raising the question of whether the results would extend to left-handed individuals, who tend to be more ambidextrous than right-handed individuals. It is worth noting however that differences in learning transfer and control have been observed between right and left handed participants in similar experiments (Wang and Sainburg, 2006).

To conclude, our results highlight differences in adaptation and behaviour between the two arms. We observed a tendency for both arms to use a more robust control strategy when facing perturbing force fields. As generally expected, the dominant arm showed better adaptation than the non-dominant arm, with the non-dominant arm relying on a more robust control strategy. Our results also suggest that both arms share a similar mechanism of adaptation.

## Acknowledgements

The authors would like to thank Tanguy de Vermont for his help in the collection of the data.

